# Infection of human organoids supports an intestinal niche for *Chlamydia trachomatis*

**DOI:** 10.1101/2024.03.25.586552

**Authors:** Pargev Hovhannisyan, Kathrin Stelzner, Markus Keicher, Kerstin Paprotka, Mastura Neyazi, Mindaugas Pauzuolis, Waled Mohammed Ali, Karthika Rajeeve, Sina Bartfeld, Thomas Rudel

## Abstract

Several reports suggest that intestinal tissue may be a natural niche for *Chlamydia trachomatis* infection and a reservoir for persistent infections in the human body. Due to the human specificity of the pathogen and the lack of suitable host models, there is limited knowledge on this topic. In our study, we modelled the course of the chlamydial infection in human primary gastrointestinal (GI) epithelial cells originating from patient-derived organoids. We show that GI cells are resistant to apical infection and *C. trachomatis* needs access to the basolateral membrane to establish an infection. Transmission electron microscopy analysis reveals the presence of both normal as well as aberrant chlamydial developmental forms in the infected cells, suggesting a possible cell-type specific nature of the infection. Furthermore, we show that the plasmid-encoded Pgp3 is an important virulence factor for the infection of human GI cells. This is the first report of *C. trachomatis* infection in human primary intestinal epithelial cells supporting a possible niche for chlamydial infection in the human intestinal tissue.

**Author summary:** Chlamydial infection has a high global prevalence and is a major health concern. Untreated infections may cause complications and lead to serious health problems, especially in women. Although the infection is usually localized to the genital tract, experiments performed in a mouse infection model as well as the accumulating clinical data suggest that the human gastrointestinal (GI) tract might represent a hidden infection niche and a source of reinfections. In our study, we used the advantages of the organoid technology to model the chlamydial infection in patient-derived primary GI epithelial cells. We were able to show that these cells are resistant to the infection, however, *Chlamydia* could utilize a basolateral entry route for efficient infection. *Chlamydia* form either normal or persistent-like developmental forms in these GI epithelial cells. We also showed the importance of the plasmid-mediated virulence in the infection of human GI cells. The results obtained in the GI infection model replicated phenotypes predicted and expected for *Chlamydia* human intestinal infection, and therefore support a role of the human GI tract as a potential niche for chlamydial infection.

## Introduction

*Chlamydia trachomatis* is a human-specific pathogen, which causes the most common bacterial sexually transmitted infections worldwide [1]. Different serovars of *C. trachomatis* have specific tissue tropism and cause diseases at different anatomical sites: serovars A-C cause eye infections, genital infections are usually associated with the serovars D-K and the more invasive serovars L1-L3 infect the lymphatic system [2, 3].

*Chlamydia* are obligate intracellular bacteria with a unique biphasic developmental cycle, during which they alternate between two morphologically and functionally different forms – elementary bodies (EBs) and reticulate bodies (RBs). *Chlamydia* possess complex and redundant mechanisms for host cell attachment and entry, which explains their ability to infect a wide range of cell types [4]. Several host cell receptors, including human integrin β1 receptor [5], epidermal growth factor receptor [6], fibroblast growth factor receptor [7] and Ephrin A2 receptor (EphA2) [8], have been found to be used by *C. trachomatis* to enter the host cell. EBs, the infectious forms of the organism, are adapted to survive in extracellular space [9]. Upon contact with the host cell, they induce their internalization and develop in a membrane bound compartment called inclusion, where they differentiate into RBs. After several rounds of replication, RBs re-differentiate into EBs, which are released from the cell to infect neighboring cells [9, 10]. Under stress conditions, RBs can enter a non-replicative but viable state called persistence, in order to survive the unfavorable conditions. They can re-enter the developmental cycle after the physiological conditions have normalized [1, 10].

The obligate intracellular lifestyle and human specificity of *C. trachomatis* limit the availability of relevant physiological host models to study the infection. Most of the knowledge about the interactions of *C. trachomatis* with the host is derived from *in vitro* experiments, which are mainly based on the use of transformed cell line models and which not always recapitulate the *in vivo* situation [11]. Since it is possible to infect mice with human *C. trachomatis* serovars under certain conditions, murine models have been widely used to study the immunopathogenesis. However, these do not always reflect the pathology of the disease observed in humans [12, 13]. In recent years, the advances in stem cell biology allowed the establishment of complex human primary cell-based host models, such as organoids, which are currently being actively used in the field of infection biology [14, 15]. Chlamydial infection has been recently successfully modelled and studied in both human and murine organoids derived from female genital tract tissues, such as human fallopian tube organoids [16], murine endometrial organoids [17] and human cervical organoids [18]. These studies have profoundly improved our understanding of chlamydial infection.

The majority of studies on *C. trachomatis*-host interactions focuses on the genital tract. There is only a limited number of studies addressing the infection at extra-genital sites. It is well known that *C. trachomatis* can infect the epithelium of the human rectum and pharynx, with a high prevalence in men who have sex with men [19]. There have also been reported cases of *C. trachomatis* DNA and antigens being detected in appendix and intestinal biopsies from patients [20, 21].

Besides human-specific pathogens, the genus *Chlamydia* contains species, which infect wild or domesticated animals [22]. Interestingly, gastrointestinal (GI) infection occurs in most animal hosts and the GI tract is a natural site of *Chlamydia* infection [23]. Some authors have proposed that *C. trachomatis* could have evolved as a commensal of the human GI tract [24] and that the human GI tract can be a site of persistent chlamydial infections and a possible reservoir of infections in the genital tract [23, 25]. Studies in mice demonstrated that following oral inoculation, the murine chlamydial pathogen *C. muridarum* crosses multiple GI barriers and establishes a long lasting non-pathological colonization in the large intestine [26]. It was also reported that Pgp3, a chlamydial plasmid-encoded virulence factor, is important for the colonization of the GI tract of mice as it helps *Chlamydia* to reach the large intestine by providing resistance against gastric acid in stomach and CD4^+^ T lymphocyte-mediated immunity in the small intestine [26].

In the present work, we investigated *C. trachomatis* infection in primary epithelial cells derived from different regions of the human GI tract using organoid-based host models. We show that *C. trachomatis* is able to infect the human GI cells from the basolateral, but not apical surface. Moreover, we demonstrate that in some cells chlamydial development is restricted and leads to the formation of aberrant bodies, hallmarks of persistent infection. We also reveal that the chlamydial plasmid and plasmid-encoded Pgp3 are important virulence factors for the infection of human GI cells as their absence leads to growth defects.

## Results

### *C. trachomatis* infects human primary GI epithelial cells

To test if *C. trachomatis* has the ability to infect human primary GI epithelial cells, we used human adult stem cell-derived organoids as a host model. They consist of primary epithelial cells organized into a single layer. In order to have each major region of the human GI tract represented in our studies, we modelled the infection in organoids derived from human corpus (stomach), jejunum (small intestine) and colon (large intestine) (Fig 1A, S1 Fig). GI organoids were dissociated into small multicellular fragments, infected with GFP-expressing *C. trachomatis* and re-suspended into fresh extracellular matrix (ECM) for the formation of new organoids. Starting from 24 hours post-infection (p.i.), we could clearly observe fluorescent *C. trachomatis* inclusions in both gastric and intestinal organoids (Fig 1B). However, in order to avoid the shredding of the 3D organoids, which disrupts the epithelial barrier integrity and to mimic the natural (apical) route of infection, we switched to a 2D setting by generating monolayers from organoids. In 2D configuration, GI cells grow in a patchy manner by forming dense small islands (S1 Fig), which gradually grow and eventually fuse forming a confluent monolayer. We infected the GI cells and HeLa cells (a common *C. trachomatis* host model) with *C. trachomatis* and observed that, in contrast to HeLa cells, chlamydial inclusions in the GI monolayers exhibited uneven distribution and were mostly located on the edges of the cell islands (Fig 1C). The infection pattern was maintained even after several rounds of chlamydial replication (4 and 8 days p.i.) and the inclusions were mostly observed on the edges of cell patches or at the fusion of two or more cell patches. We also observed that the infection progresses from the margins inward to the centre of the patches, presumably via successive infection of new adjacent cells. While HeLa cells were almost completely lysed by *C. trachomatis* infection on day 4, primary GI cells, especially those in the centre of the monolayers, were still intact.

**Fig 1.**
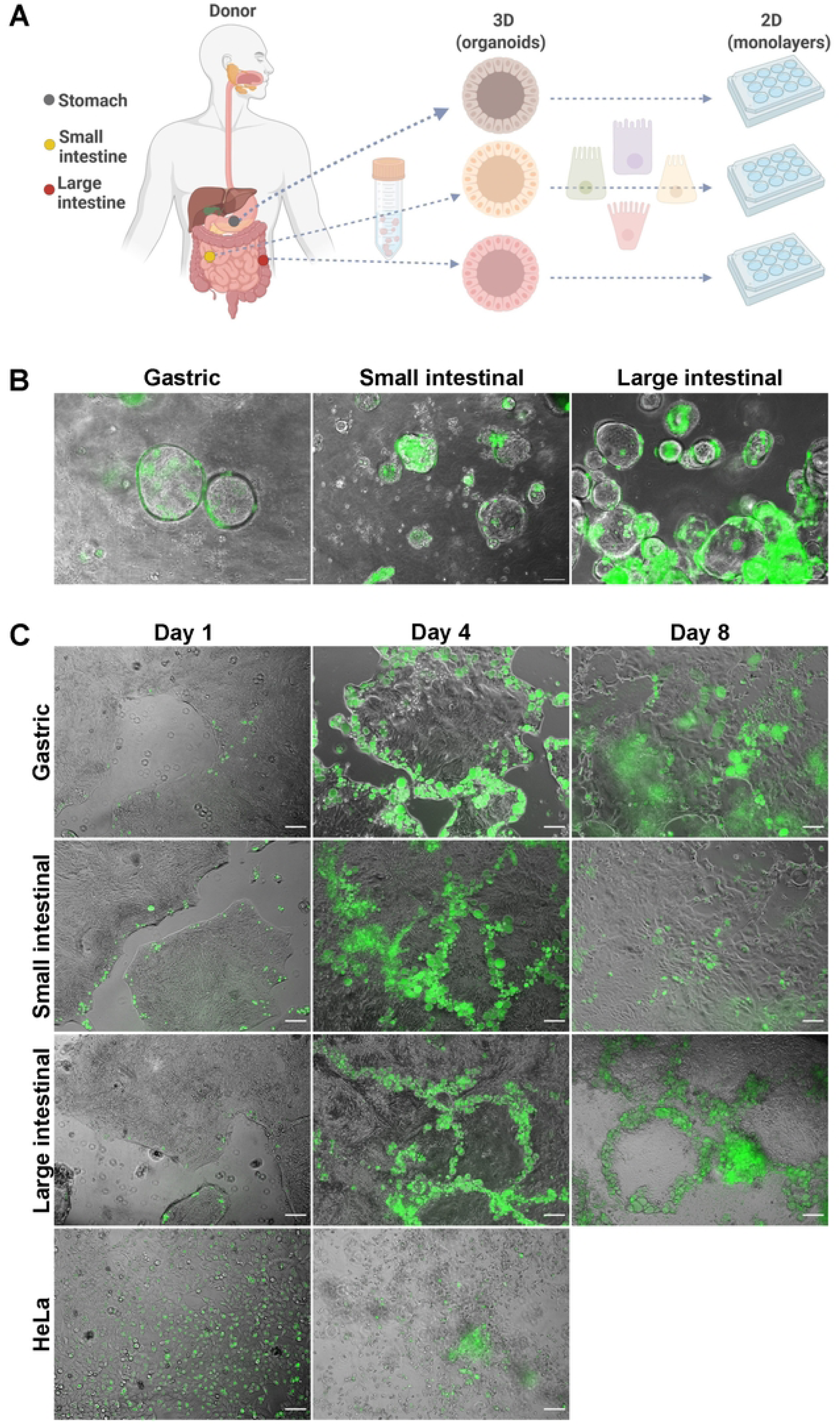
*C. trachomatis* infects patient-derived GI cells. (A) Schematic representation of host model generation. (B) Human gastric and intestinal organoids infected with GFP-expressing *C. trachomatis* (MOI of 5) at 48 hours p.i. (C) Organoid-derived subconfluent monolayers of human primary gastrointestinal epithelial cells and HeLa cells were infected with GFP-expressing *C. trachomatis* (MOI of 0.5) and observed daily. Shown are representative images obtained at 1, 4 and 8 days p.i.. Images in (B) and (C) were taken with phase-contrast and in the green fluorescence channel and merged. Scale bar: 100 µm.

### Disruption of the cell junctions increases the efficiency of *C. trachomatis* infection in GI cells

The observed novel pattern of *C. trachomatis* infection in GI monolayers could either be attributable to the physical properties of cells in monolayers or be a consequence of cell-type/state specific infection events. As we observed that proliferative cells are often found closer to the edges of cell islands in GI monolayers, we asked whether *Chlamydia* might predominantly infect actively dividing cells. We co-stained the infected gastric monolayers for Ki67, which is a nuclear protein and a marker of proliferation, and determined the number of inclusions associated with Ki67-positive or negative cells (S2A, S2B Fig). We could not find a direct link between the cell proliferation status and the infection, as the microscopic analysis revealed a similar infection burden in Ki67-positive and negative cells.

We also observed that, in contrast to HeLa cells, GI monolayer cells have a dense arrangement within individual cell patches and hypothesized that *Chlamydia* might preferentially infect the edges of the patches due to the distinct localization of the entry receptors on the basolateral surface, e.g. at cell-cell junctions, and their absence on the apical surface of the cells. To test this hypothesis, we disrupted the cell-cell junctions by pre-treating the gastric cells with the Ca^2+^ chelating agent ethylene glycol-bis(beta-aminoethyl ether)-N,N,N’,N’-tetraacetic acid (EGTA), which was previously reported to cause disruption of tight junctions and increase the paracellular space [27, 28]. We applied live cell imaging to monitor the effect of EGTA on primary gastric and HeLa cells.

Characteristic morphological changes, such as gradually increasing distance between neighbouring cells and loss of monolayer integrity, were observed in the treated gastric cells, whereas HeLa cells exhibited a more defined and elongated shape (Fig 2A). Following EGTA pre-treatment, gastric and HeLa cells were infected with *C. trachomatis* and subjected to microscopy analysis 24 hours p.i.. It revealed a more dispersed infection pattern in the EGTA-treated gastric cells compared to untreated control cells (Fig 2B), as well as a significantly higher infection rate in gastric cells pre-treated with EGTA for 30 min (approx. 1.7-fold higher) or 60 min (approx. 2.8-fold higher) (Fig 2C). We detected no changes in the infection pattern or rate of HeLa cells upon pre-treatment with EGTA (Fig 2B, 2C). The cell morphology and the infection pattern of intestinal cells were affected in a similar manner upon EGTA pre-treatment (S3A, S3B Fig).

**Fig 2.**
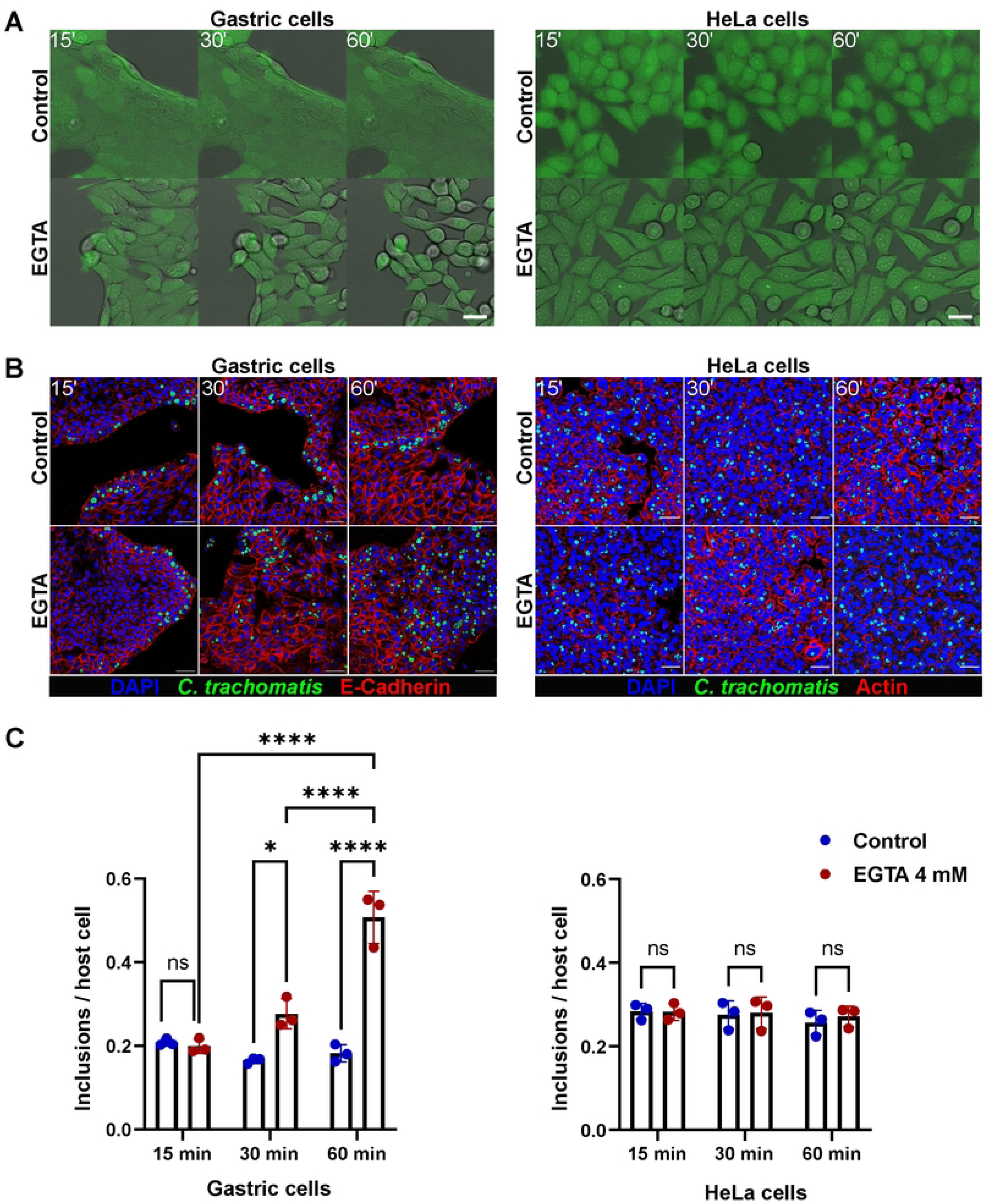
Disruption of the cell junctions affects the infection rate and pattern in gastric cells. (A) Stills from live cell imaging experiment performed to monitor the effect of 4 mM EGTA on the morphology of human gastric (left panel) and HeLa (right panel) cells over the time course of treatment (green: CellTracker dye, gray: brightfield). Scale bar: 25 µm. (B) Representative confocal fluorescence images showing the effect of EGTA treatment on the outcome of infection in gastric (left panel) and HeLa (right panel) cells. Gastric and HeLa cells were pre-treated with 4 mM EGTA (for 15, 30 or 60 min) or left untreated, and infected with GFP-expressing *C. trachomatis* (MOI of 5 and 0.5, respectively). 24 hours p.i. the cells were fixed, stained and subjected to confocal microscopy (blue: DAPI, green: *C. trachomatis*, red: E-Cadherin or Actin). Scale bar: 50 µm. (C) To measure the changes of the infection rate caused by EGTA pre-treatment, the numbers of chlamydial inclusions and host cell nuclei were determined in 14 fields of view per sample by automated microscopy. Data represent mean values ± SD from three independent experiments. Statistical analysis was performed using two-way ANOVA (ns = not significant, *P<0.05, ****P<0.0001).

### *C. trachomatis* infects the GI cells via the basolateral route

As GI cells with disrupted junctions could be efficiently infected by *C. trachomatis*, we next tested the ability of the pathogen to selectively infect gastric and intestinal monolayers from the apical versus the basolateral surface. Culturing the cells on a porous membrane in cell culture inserts allowed a separation of these two surfaces. Gastric and small intestinal primary cells were seeded on the membrane and grown until they formed a confluent monolayer with a polarized phenotype, where the basolateral side of the cells was oriented to the cell culture insert membrane. The cells were infected with *C. trachomatis* from the apical or basolateral side (Fig 3A) and analyzed 24 hours p.i. by confocal microscopy. Interestingly, we detected no or extremely few inclusions in apically infected gastric and intestinal monolayers, whereas the basolateral infection was highly efficient (Fig 3B). To check whether the observed phenotype of basolateral infection is specific to human GI cells only or might also be common for other columnar epithelial cell types, we performed the infection assay in organoid-derived human primary fallopian tube epithelial cells (S4A Fig, S4B Fig). Although we detected inclusions in apically infected fallopian tube samples, the basolateral infection resulted in a significantly higher infection rate.

**Fig 3.**
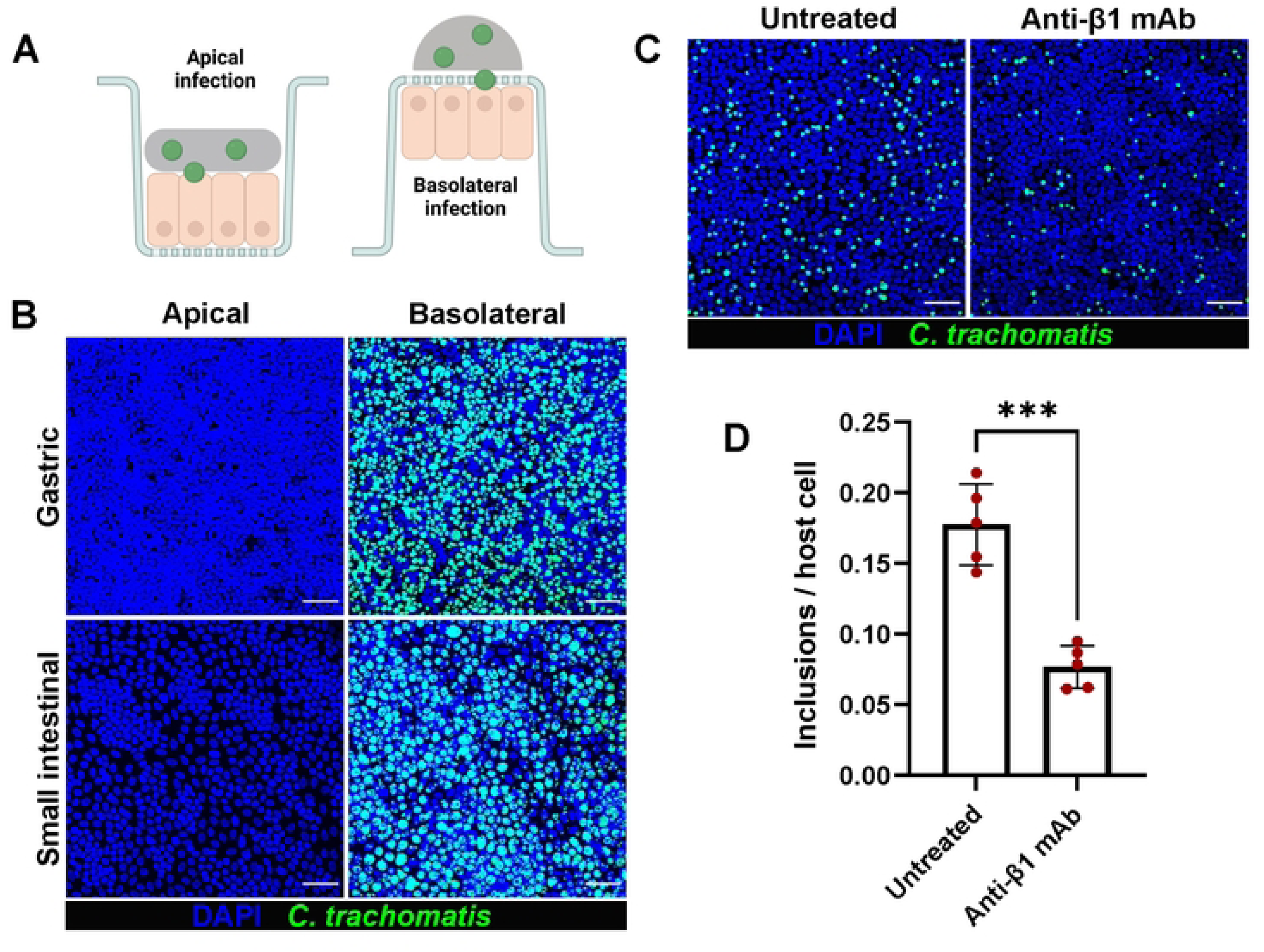
*C. trachomatis* infects human GI cells from the basolateral surface. (A) Schematic representation of the infection setup. (B) Gastric and small intestinal cells grown on cell culture inserts were infected with GFP-expressing *C. trachomatis* (MOI of 5) from the apical or basolateral surface and after 2 hours of incubation the inoculum was removed, and the cells were kept in fresh organoid medium. 24 hours p.i. the cells were fixed, stained and subjected to confocal microscopy. Shown are representative confocal fluorescence Z-stack images of at least three independent experiments (blue: DAPI, green: *C. trachomatis*). Scale bar: 50 µm. (C) The gastric cells grown on cell culture inserts were pre-treated with 25 µg/ml blocking anti-integrin β1 antibodies for 1 hour or left untreated, infected with GFP-expressing *C. trachomatis* (MOI of 5) for 2 hours in the presence of blocking antibodies. 24 hours p.i. cells were fixed, stained and subjected to confocal microscopy. Shown are representative confocal fluorescence Z-stack images of five independent experiments (blue: DAPI, green: *C. trachomatis*). Scale bar: 50 µm. (D) The numbers of chlamydial inclusions and host cell nuclei in (C) were quantified in at least 10 fields of view per sample using Fiji. Data represent mean values ± SD from five independent experiments. Statistical analysis was performed using an unpaired t-test (***P<0.001).

Next, we aimed to identify a basolaterally expressed host cell receptor, which could potentially be involved in the invasion of GI cells by *C. trachomatis*. Integrin receptors, including β1-integrins, are usually located at the basolateral surface of polarized epithelial cells [29, 30]. Integrin β1 receptor is also exploited by *C. trachomatis* for host cell entry among several other receptors [5]. To determine its involvement in the infection, we treated the gastric cells grown on cell culture inserts with anti-integrin β1 blocking antibodies and infected with *C. trachomatis*. Confocal microscopy analysis of the infected samples 24 hours p.i. revealed a significantly reduced (2.3-fold) infection rate in antibody-treated cells compared to untreated cells (Fig 3C, 3D). Collectively, our findings indicate that human GI epithelial cells are highly resistant to *C. trachomatis* apical infection and an access to the receptors localized on the basolateral surface of the cells is needed for efficient infection.

### Chlamydial plasmid-encoded Pgp3 is important for propagation in human GI cells

The chlamydial plasmid, particularly the plasmid encoded virulence factor Pgp3, is important for the colonization of the GI tract of the mice by *C*. *muridarum* [31]. Pgp3 has been found to play a critical role in protecting *Chlamydia* against gastric acid killing and thus allowing its further dissemination into the lower GI tract of the mice. We asked whether chlamydial plasmid and Pgp3 are involved in the infection of human GI cells and therefore compared the infectivity of wild-type (*Ctr* WT), Pgp3-deficient (*Ctr* Δ*pgp3*) and plasmid-free (*Ctr* PF) *C. trachomatis* strains in human gastric, small and large intestinal epithelial cells (Fig 4A). We first titrated the infectivity of the strains in HeLa cells to reach similar infection load in order to enable a cross-strain comparison in GI cells and obtained a similar infection rate for *Ctr* WT and *Ctr* Δ*pgp3* and a slightly higher rate for *Ctr* PF (S5A Fig). Next, we performed the infectivity assay in human GI cells derived from three different donors using proportionate bacterial loads. During primary infection in gastric cells, *Ctr* Δ*pgp3* and *Ctr* PF showed significantly reduced infectivity (approx. 2.3-fold and 2.9-fold lower, respectively) compared to *Ctr* WT (Fig 4B) and a similar trend was observed in small and large intestinal cells (Fig 4C, 4D). To assess chlamydial development and fitness, equal amounts of cell lysates were transferred onto freshly seeded HeLa cells 48 hours p.i. and analyzed 24 hours p.i. (here referred to as progeny infection). *Ctr* Δ*pgp3* from gastric cells formed significantly fewer (2.8-fold fewer) infective progeny compared to *Ctr* WT (Fig 4E) and a similar trend was observed in intestinal cells (Fig 4F, 4G). *Ctr* WT and *Ctr* Δ*pgp3* grown in HeLa cells formed equal amount of progeny, and *Ctr* PF formed significantly fewer (approx. 1.8-fold fewer) progeny compared to *Ctr* WT and *Ctr* Δ*pgp3* despite the initially higher load in primary infection (S5B Fig).

**Fig 4.**
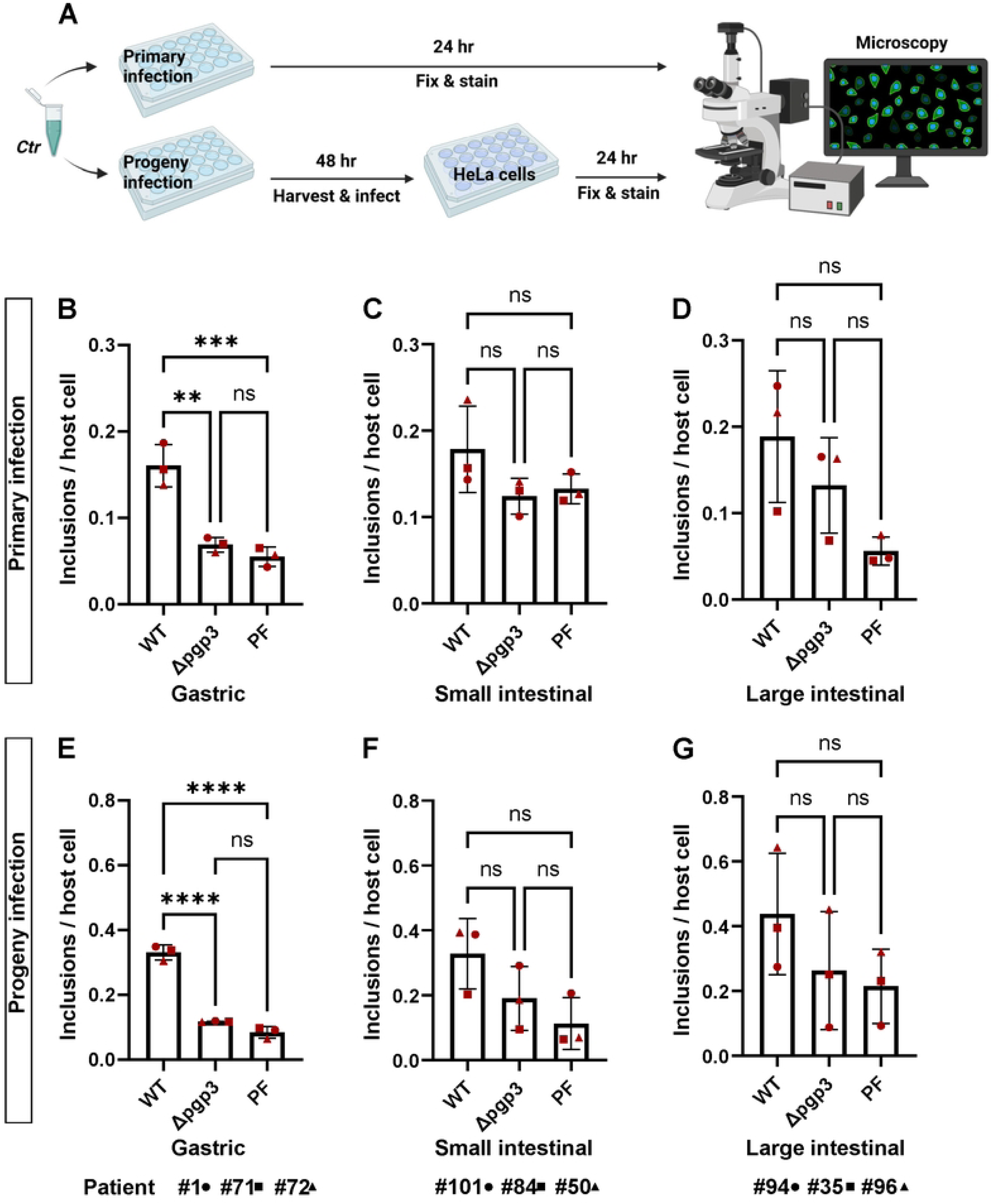
Pgp3-deficient *C. trachomatis* shows a growth defect in human GI cells. (A) Schematic representation of the primary and progeny infection assays. To compare the infectivity of *Ctr* WT, *Ctr* Δ*pgp3* and *Ctr* PF strains in gastric (B), small intestinal (C) and large intestinal (D) cells, subconfluent monolayers of the cells were infected using MOI of 5. 24 hours p.i. the cells were fixed, stained and the infection rate was determined by quantifying the numbers of inclusions and host cell nuclei in 14 fields of view per sample by automated microscopy. To assess the infectivity of the chlamydial progeny, the infected cells were lysed 48 hours p.i. and freshly seeded HeLa cells were infected with an aliquot of the lysates. 24 hours p.i. the HeLa cells were fixed, stained and the infection rates for the chlamydial progenies from gastric (E), small intestinal (F) and large intestinal (G) cells were determined in 14 fields of view per sample by automated microscopy. All graphs represent mean values ± SD from three independent experiments. Statistical significance was determined by one-way ANOVA (ns = not significant, **P<0.01, ***P<0.001, ****P<0.0001). The data point numbers and shapes below the graphs refer to the donor IDs used in the experiments.

To confirm the results by measuring another parameter for chlamydial development, we also determined the average size of the inclusions during primary infection. Interestingly, although *Ctr* WT and *Ctr* Δ*pgp3* formed inclusions of similar size in HeLa cells (S5C Fig), in gastric cells *Ctr* Δ*pgp3* formed significantly smaller inclusions compared to *Ctr* WT (Fig 5A) and a similar trend was observed in intestinal cells (Fig 5B, 5C). Taken together, these results suggest that the lack of the Pgp3 reduces the infectivity of *C. trachomatis* in human primary gastric and possibly also in intestinal epithelial cells.

**Fig 5.**
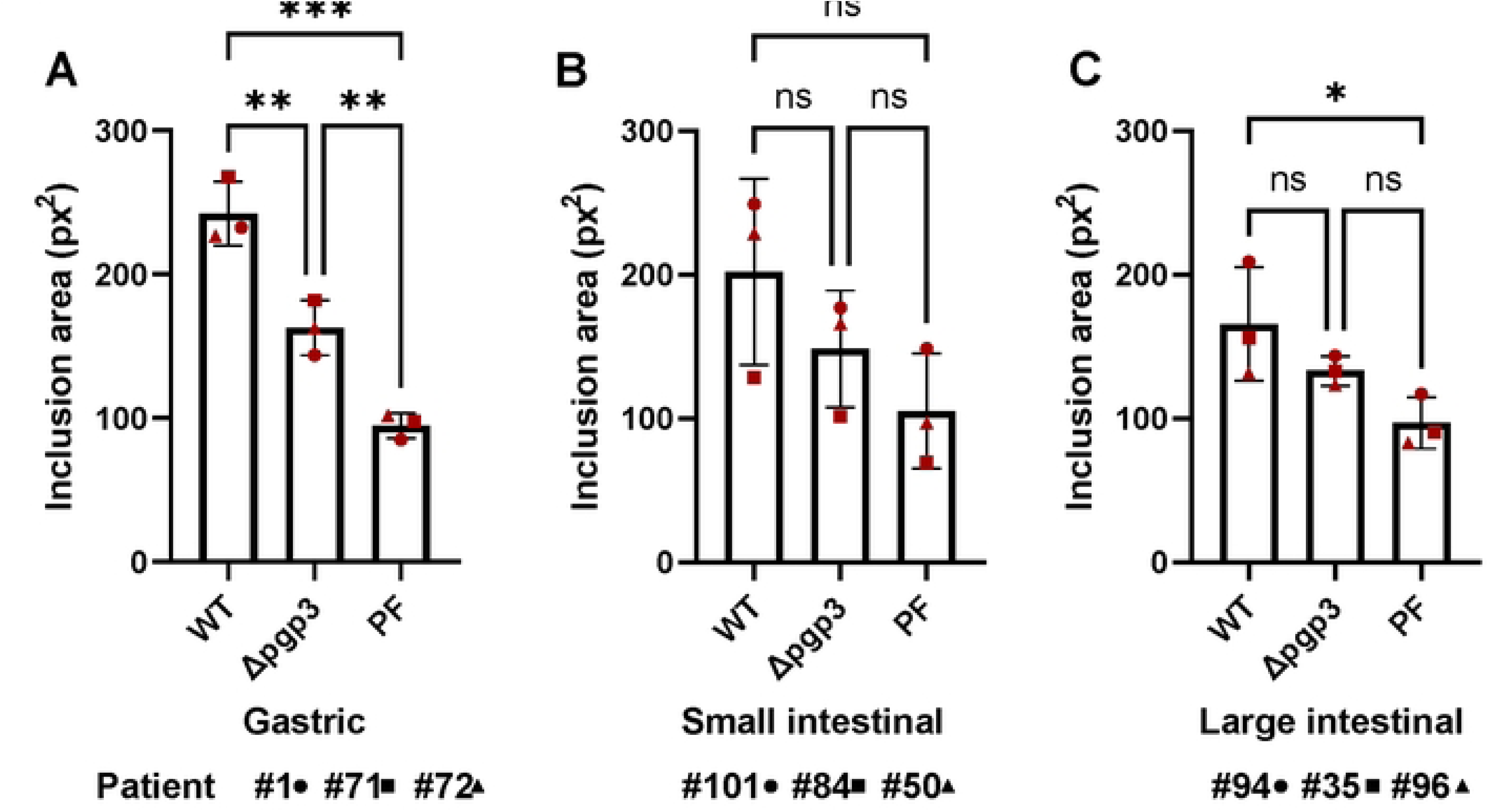
Pgp3-deficient *C. trachomatis* forms smaller inclusions in human GI cells. Subconfluent monolayers of gastric (A), small intestinal (B) and large intestinal (C) cells were infected with *Ctr* WT, *Ctr* Δ*pgp3* and *Ctr* PF strains at MOI of 5. 24 hours p.i. the cells were fixed, stained and the average size of the inclusions was determined by automated microscopy in 14 fields of view per sample. All graphs represent mean values ± SD from three independent experiments. Statistical significance was determined by one-way ANOVA (ns = not significant, *P<0.05, **P<0.01, ***P<0.001). The data point numbers and shapes below the graphs refer to the donor IDs used in the experiments.

### Human GI cells harbour aberrant developmental forms of *C. trachomatis*

Human intestinal tissue is hypothesized to represent a potential niche for persistent chlamydial infection, which is frequently characterized by the appearance of so-called aberrant bodies (ABs), irregular particles much larger than RBs [23, 32]. We therefore characterized chlamydial inclusions in human GI cells at the ultrastructural level by using transmission electron microscopy (TEM). Gastric and intestinal cells derived from 3 different donors were infected with *C. trachomatis* and 40 hours p.i. processed for TEM. Surprisingly, we found a mixed population of normal (Fig 6A) and aberrant (Fig 6B) developmental forms of inclusions in both gastric and intestinal cells. Morphological evaluation of the obtained micrographs revealed different morphologies of aberrant inclusions, some of them containing exclusively enlarged ABs and some others harbouring also dividing forms of RBs or even EBs. In contrast to GI cells, no ABs could be detected in infected HeLa cells. This indicates that gastric and intestinal cells could harbour ABs and thus potentially may serve as a reservoir for persistent infection.

**Fig 6.**
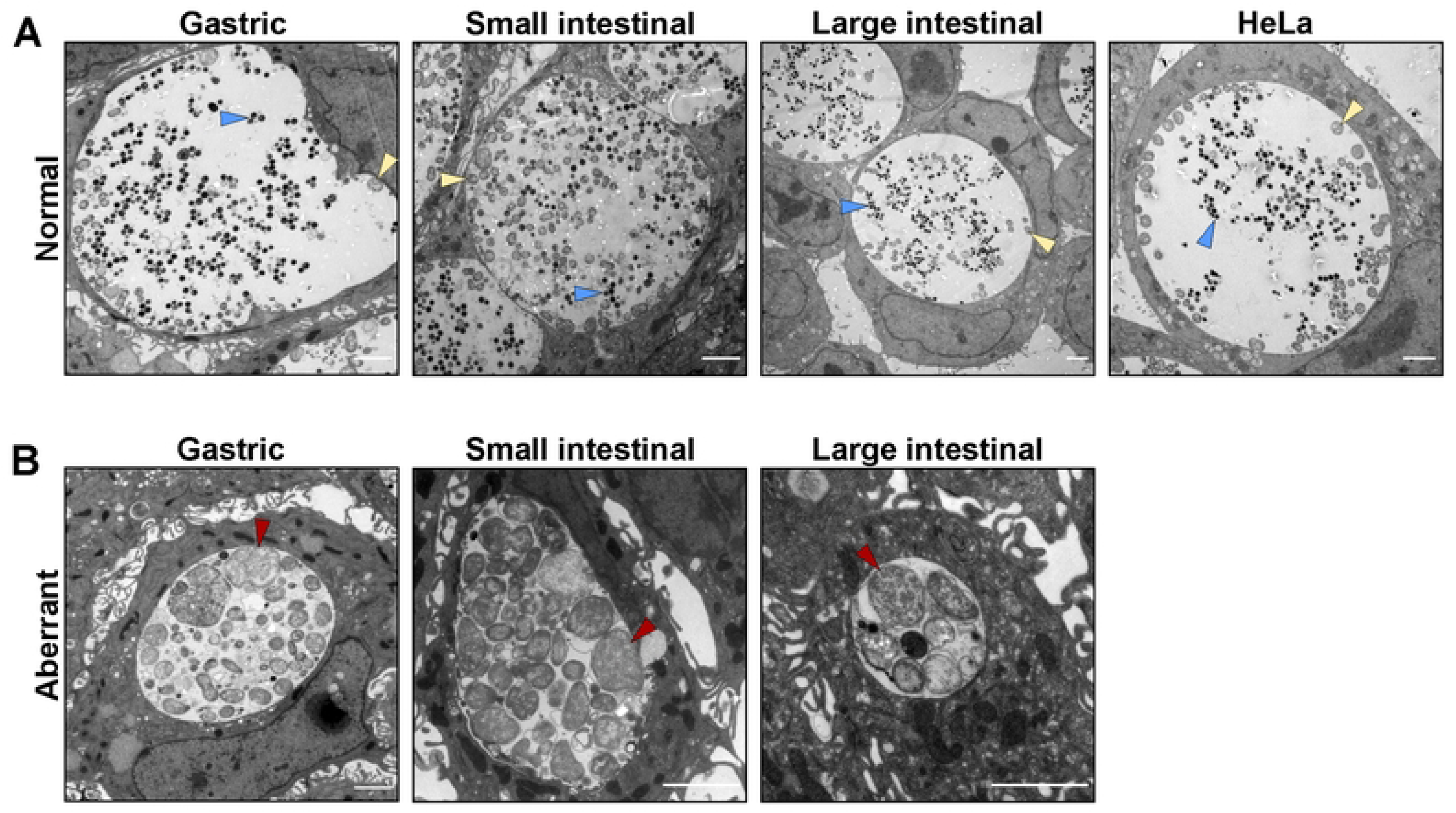
Human GI cells harbour aberrant inclusions. Subconfluent monolayers of gastric, small and large intestinal epithelial cells derived from three different donors were pre-treated with EGTA and infected with *C. trachomatis* at MOI of 5 and HeLa cells were infected at MOI of 0.5. 40 hours p.i. the cells were fixed and processed for TEM. Shown are representative images of the normal (A) and aberrant-like (B) inclusions found in the respective cells. Triangles indicate EBs (blue), RBs (yellow) and ABs (red). Scale bar: 2 µm.

## Discussion

Increasing evidence suggests that the mucosa of the GI tract provides a niche for persistent *C. trachomatis* infections in the human body and can potentially cause repeated infections in other tissues, including the genital tract [23]. Nevertheless, there is only a limited number of studies on the pathogenesis of *C. trachomatis* in human GI cells and most of the knowledge comes from murine infection models. So far, *C. trachomatis* GI infection has been studied using cancer cell line models, such as human enteroendocrine LCC-18 and CNDT-2 cells [33], Caco-2 and COLO-205 colon carcinoma cells [34]. In the present study, we modelled *C. trachomatis* infection in human primary gastric and intestinal epithelial cells.

Our results based on 3D and 2D infection models indicate that both gastric and intestinal cells can support the chlamydial development. However, attachment of *Chlamydia* to the basolateral membrane is necessary for establishing an infection. In the GI monolayer model, the tightly arranged cell architecture renders the cells resistant to infection from the apical surface, whereas the infection from the basolateral surface is highly efficient (Fig 3B). In line with this, in subconfluent GI monolayers *C. trachomatis* infects the marginally located cells, even after completion of several chlamydial developmental cycles (Fig 1C). We assume this is due to the localization of the chlamydial entry receptors on the basolateral, but not on the apical surface of the cells. Consequently, only the cells on the edges of the cell patches allow *Chlamydia* to access the basolateral side, while other cells are engaged in junctional complexes. This theory is substantiated by the observation that disruption of the cell-cell contacts by the Ca^2+^ chelating agent EGTA increases the number of infected cells and renders the infection pattern more random (Fig 2B, 2C). In polarized epithelial cells tight and adherens junctions shield the basolaterally expressed integrins [29], which prompted us to assess the role of the known chlamydial entry receptor integrin β1 as a potential candidate receptor. Blocking of the receptor on the basolateral surface of the gastric cells led to a decreased infection rate, confirming its involvement in the infection (Fig 3C, 3D). Recently, it has been demonstrated that EphA2, another chlamydial entry receptor, is strictly localized at cell-cell junctions in primary gastric epithelial cells [15]. Thus, further studies are needed to evaluate the involvement of EphA2 and other basolaterally positioned receptors in the *C. trachomatis* infection of human GI cells.

To check if the basolateral route of chlamydial infection might take place in other columnar epithelial cell models as well, we modelled the infection in human primary fallopian tube cells. Although we could detect inclusions in apically infected samples, the basolateral infection resulted in a significantly higher infection rate (S4A, S4B). This might explain the low infection rate (<5%) previously reported in a similar apically infected fallopian tube model [35]. However, it is to note that in contrast to GI cells, the culture of human fallopian tube cells on 3 µm pore-size membranes (a technical requirement imposed by the experimental setup) compromises their ability to polarize and hinders the comparison with the GI model.

The importance of the plasmid for the successful colonization of the GI tract by *Chlamydia* has been demonstrated in the mouse model of infection [26]. The importance of the plasmid is further evidenced by the fact that naturally occurring plasmid-free clinical isolates are very rare [36, 37].

Following oral inoculation, *Chlamydia* overcome several GI barriers in mice in order to reach and colonize the large intestine, and plasmid lacking mutants, in particular Pgp3-mutants, are defective in colonizing the upper GI tract of the mice [26, 31]. It is speculated that *Chlamydia* use plasmid-encoded factors to colonize the upper GI tract, whereas chromosome-encoded factors are more important for colonization of the lower GI tract [26]. We compared the infectivity of the wild-type, Pgp3-deficient and plasmid-free *C. trachomatis* in human GI cells to check if the plasmid-associated infectivity defects reported *in vivo* could be reproduced in a human system *in vitro*. Compared to the wild-type, *Ctr* Δ*pgp3* formed significantly fewer and smaller inclusions in human gastric cells and a similar trend was observed in intestinal cells (Fig 4, Fig 5). In mice Pgp3 plays a role in resisting gastric acid in stomach and evasion of the CD4+ T lymphocyte-mediated immunity in the small intestine [26]. However, our epithelial model does not contain immune cells nor is gastric acid produced because the appropriate cells (parietal cells) are absent in the system. Further studies are needed to clarify the specific role of Pgp3 during the infection of GI epithelial cells.

In search of signs of persistence, we performed electron microscopic analysis of infected GI cells, which revealed that gastric, small and large intestinal cells infected with *C. trachomatis* harbor not only normal developed inclusions, but also aberrant developmental forms, which have different morphologies (Fig 6). 2D monolayer cultures of primary GI cells are not monotypic and contain different cell types and we assume that possibly some cell types restrict, while some others permit chlamydial development. Interestingly, inclusions of similar morphology have been found in the intestine of the pigs naturally or experimentally infected with *C. suis* [38].

Taken together, our infection model replicated phenotypes predicted and expected for *C. trachomatis* human intestinal infection, like the occurrence of persistent infection, and therefore supported a role of the human GI tract as a potential niche for chlamydial infection. At the same time, our results in 2D monolayers imply that the healthy intact GI epithelium is resistant to luminal *C. trachomatis* infection. It is likely that prior events affecting the epithelial barrier integrity and polarity, such as inflammation, epithelial-mesenchymal transition or mechanical injuries, could be necessary for the establishment of infection. Our study also highlights the importance of using physiologically relevant host models for modelling host-pathogen interactions. There are, however, clear limitations of our model like the absence of natural microbiota and a functional innate and adaptive immune system. The intestinal microbiota and immune system are important factors that protect the intestinal epithelium from pathogens [39]. Initial attempts have been undertaken to generate human intestinal models complemented with human microbiota and innate immune cells [40, 41]. Although these systems still do not adequately resemble the natural human intestinal environment, they will be useful to further investigate the role of chlamydial infection in the human GI tract.

## Methods

### Ethics statement

This study was approved by the ethical committee of the University Hospital of Würzburg (approval 37/16 and 36/16).

### Culture of organoids

Patient-derived gastric (corporal), small intestinal (jejunal), large intestinal (colonic) and fallopian tube organoid lines were retrieved from an established biobank and cultured in Matrigel drops (Corning, 356231) overlaid with the corresponding organoid culture medium in 24-well plates and kept in a humidified incubator at 37°C and 5% CO_2_. Advanced DMEM/F12 (Thermo Fisher Scientific, 12634028) supplemented with 10 mM HEPES (Thermo Fisher Scientific, 15630056) and 1% GlutaMAX (Thermo Fisher Scientific, 35050061) was used as a basal medium and the organoid-specific growth factors were added to it (full composition of the media is provided in S1 Table).

Organoids were passaged every 7-14 days at a ratio 1:2 to 1:10 depending on the donor and the medium was changed every 2-3 days. During the first two days of culture after splitting, media were supplemented with 10 µM ROCK inhibitor (AbMole Bioscience, Y-27632).

### Culture of organoid-derived 2D monolayers

2D monolayers in the microwell plates were generated according to previously established protocols [42] (Neyazi et al., in preparation): organoids were dissociated into single cells using TrypLE (Thermo Fisher Scientific, 12604013) at 37°C, seeded into the microwell plate and grown in corresponding organoid culture medium at 37°C and 5% CO_2_. To generate monolayers on cell culture inserts (Sigma Aldrich, PITP01250), dissociated cells were seeded onto the membrane and grown in corresponding organoid culture medium till reaching confluency at 37°C and 5% CO_2_.

### Culture of cell lines

Human cervix adenocarcinoma cells (HeLa 229, ATCC CCL-2.1^TM^) were cultured in RPMI-1640 medium (Thermo Fisher Scientific, 72400054) supplemented with 10% fetal calf serum (FCS) (Sigma Aldrich, F7524) and maintained in a humidified incubator at 37°C and 5% CO_2_.

### *C. trachomatis* strains and cultivation

All *C. trachomatis* strains used in the study originate from the *C. trachomatis* L2 (434/Bu, ATCC VR-902B). The GFP-expressing strain was generated by transforming *C. trachomatis* L2 with pGFP::SW2 plasmid as previously described [43]. The plasmid-free strain was generated using novobiocin treatment as previously described [36]. The Pgp3 deletion mutant was generated by transforming the plasmid-free *C. trachomatis* L2 with pGFP::SW2Δpgp3 plasmid.

For stock preparation, the strains were propagated in HeLa 229 cells for 48 hours, after which the cells were lysed with glass beads and *Chlamydia* were separated by centrifugation at 2,000 x g for 10 min at 4°C. *Chlamydia* were afterwards pelleted by centrifugation at 30,000 x g for 30 min at 4°C and washed with 1x sucrose-phosphate-glutamic acid buffer (SPG). The bacterial pellet was re-suspended in 1x SPG buffer, aliquoted and stored at -80°C. For infectivity titration, HeLa 229 cells grown in 24-well microplates (Ibidi, 82426) were infected with different amounts of bacteria for 2 hours and 24 hours p.i. the cells were fixed and stained (details of the staining provided in Immunofluorescence). The number of chlamydial inclusions and host cell nuclei was measured and analyzed with Operetta automated microscopy system (Perkin-Elmer). The amount of the *Chlamydia* resulting in a *Chlamydia* inclusion to HeLa cell nuclei ratio of 0.5 was determined and the obtained infection rate was considered as a multiplicity of infection (MOI) of 0.5. Ten times more bacteria were used for the infection of primary epithelial cells (MOI relative to HeLa cells).

### Infection of the organoids

Organoids were infected following a published protocol [16]: briefly, mature organoids were released from Matrigel drops with cold basal medium and mechanically disrupted into small fragments. Fragmented organoids were centrifuged (300 x g for 5 min at 4°C) and the pellet was infected with GFP-expressing *C. trachomatis* L2 for 20 min at MOI of 5. After incubation, the pellet was re-suspended in fresh Matrigel and seeded in a microwell plate. The infection was monitored daily using phase-contrast and fluorescence microscopy (Leica DMI3000B).

### Infection of the monolayers

For the 2D infections in microwell plates, HeLa and primary GI cells were seeded in 24-well microplates (Ibidi, 82426) and grown to 70% confluency. For the 2D long-term infections (Fig 1C), HeLa and GI cells were infected with *C. trachomatis* at MOI of 0.5 and after 2 hours of incubation, the medium of the plates was exchanged. Images were obtained with a phase-contrast and fluorescence microscope (Leica DMI3000B) every 24 hours and the medium change was performed as needed. For the 2D infectivity assays HeLa cells were infected at MOI of 0.5 and the primary GI cells were infected at MOI of 5. After 2 hours of infection, the medium of the plates was exchanged. To assess the primary infection, 24 hours p.i. the cells were fixed, stained and the numbers of inclusions and the host cell nuclei were quantified with Operetta automated microscopy system. To assess the infectivity (progeny infection), infected cells were lysed with glass beads 48 hours p.i. and dilutions of the supernatant were used to infect freshly seeded HeLa cells, which were fixed 24 hours p.i., stained and analyzed with Operetta automated microscopy system.

For infection assays in cell culture inserts, the bacterial inoculum resuspended in basal medium to a volume of 75 µl, was applied to the apical surface of cells in cell culture inserts or carefully added as a drop to the basolateral side on inverted inserts. After 2 hours of incubation, the inoculum was removed and the inserts were kept in the corresponding growth media. For receptor-blocking assays, the basolateral surfaces of the cells were pre-treated with blocking anti-integrin β1 antibody (25 µg/ml, R&D Systems, MAB17781) for 1h at 37°C, followed by infection with *C. trachomatis* L2 at MOI of 0.5 in the presence of antibodies. After 2 hours of infection, the inoculum was removed and the cells were kept in corresponding growth media. The cells were fixed 24 hours p.i. and analyzed via confocoal microscopy. The numbers of chlamydial inclusions and host cell nuclei were determined using Fiji.

To induce disruption of the cell-cell junctions, cells were treated with 4 mM EGTA (Sigma Aldrich, E4378) for the specified amount of time at 37°C prior to infection.

### Immunofluorescence

At designated times, cells were washed with phosphate buffered saline (PBS, Thermo Fisher Scientific, 14190169) and fixed with 4% paraformaldehyde (PFA, Morphisto, 11762.01000). After permeabilization with 0.2% Triton X-100 (Roth, 3051.4) in PBS, cells were blocked with 2% FCS or 1% bovine serum albumin (Roth, 8076.3) in PBS. Following primary antibodies were used: anti-HSP60 (Santa Cruz, sc-57840), anti-E-cadherin (Becton Dickinson, 560064), anti-Ki67 (Cell Signaling, 9129S). Alexa Fluor (Thermo Fisher Scientific) or Cy (Dianova) conjugated secondary antibodies were used. Actin filaments were stained with Phalloidin (MoBiTec, MFP-D555-33) and the DNA with DAPI (Sigma Aldrich, D9542). Images were collected using a confocal microscope (Leica, TCS SP5) or Operetta automated microscopy system (Perkin-Elmer).

### Live cell imaging

Gastric and HeLa cells were seeded in 8-well chamber µ-slides (ibidi, 80826) and grown in corresponding media untill reaching 70% confluency. Prior to imaging cells were incubated with 2.5 µM CellTracker™ Green (Thermo Fisher Scientific/Invitrogen, C2925) fluorescent probe for 30 min at 37°C and subsequently washed with PBS. Time-lapse imaging was performed on a Leica TCS SP5 confocal microscope with a 40x oil immersion objective (Leica HC PL APO CS2, NA=1.3) 5 min after addition of 4 mM EGTA. During the imaging cells were incubated at 37°C and 5% CO_2_ using a live-cell incubation chamber (Life Imaging Systems) and images were recorded in 1 min time intervals in 8-bit mode at a resolution of 1024×1024. The LAS AF software (Leica) was used for setting adjustment and image acquisition, and Fiji for image-processing [44].

### Transmission electron microscopy

Primary GI and HeLa cells were cultured in 24-well microplates (Ibidi, 82426) as subconfluent monolayers and infected with wild-type *C. trachomatis* L2 at MOI of 5 and 0.5, respectively. Prior to infection, the GI cells were pre-treated with 4 mM EGTA for 30 min. 40 hours p.i., cells were washed with PBS, fixed with 2.5% glutaraldehyde (Sigma) for 30 min at room temperature, washed with 50 mM Cacodylate buffer (Roth), incubated in OsO_4_/Cacodylate buffer for 1 hour and in 0.5% Uranylacetat overnight. Dehydrated samples were embedded in EPON and cut. Images were taken with JEOL JEM-1400 Flash microscope.

### Statistical analysis

The experimental data were analyzed using Graphpad Prism 9.3.1 software. For the analysis at least three independent experiments were used, unless otherwise indicated. Statistical significance was determined using unpaired Student’s t-test between two groups and one-way or two-way ANOVA between multiple groups. Data are shown as mean ± SD. A P value of less than 0.05 represented a statistically significant difference.

## Acknowledgements

We thank Claudia Gehrig-Höhn and Daniela Bunsen for their support in preparation of samples for TEM and imaging; Dr. Carmen Aguilar and Dr. Özge Kayisoglu for valuable advice. Figures 1A, 3A and 4A were made in Biorender.com.

## Supporting information

**S1 Table.** Media composition for human gastric, intestinal and fallopian tube organoids (CM: conditioned medium; EGF: epidermal growth factor; FGF-10: fibroblast growth factor-10; TGF-β: transforming growth factor-β; IGF-1: insulin-like growth factor I; FGF-2: fibroblast growth factor-basic)

| Human gastric organoid medium |  |  |  |
| --- | --- | --- | --- |
| Reagent | Supplier | Catalog Number | Final concentration |
| Basal medium |  |  | 30% |
| WNT CM | stable cell line |  | 50% |
| R-Spondin CM | stable cell line |  | 10% |
| Noggin CM | stable cell line |  | 10% |
| B27 supplement | Thermo Fisher Scientific | 12587010 | 1x |
| N-acetylcysteine | Sigma Aldrich | A9165 | 1.25 mM |
| EGF | Peptrotech | AF-100-15 | 50 ng/mL |
| FGF-10 | Peptrotech | 100-26 | 100 ng/mL |
| Gastrin-I | Tocris | 3006 | 1 nM |
| TGF-β inhibitor | Tocris | 2939 | 2 uM |
| Human intestinal organoid medium |  |  |  |
| Reagent | Supplier | Catalog Number | Final concentration |
| Basal medium |  |  | 30% |
| WNT CM | stable cell line |  | 50% |
| R-Spondin CM | stable cell line |  | 10% |
| Noggin CM | stable cell line |  | 10% |
| B27 supplement | Thermo Fisher Scientific | 12587010 | 1x |
| N-Ac | Sigma Aldrich | A9165 | 1.25 mM |
| EGF | Peptrotech | AF-100-15 | 50 ng/mL |
| IGF-1 | Biolegend | 590904 | 100 ng/mL |
| FGF-2 | Peptrotech | AF-100-18B | 50 ng/mL |
| Gastrin-I | Tocris | 3006 | 10 nM |
| TGF-β inhibitor | Tocris | 2939 | 0.5 uM |
| Human fallopian tube organoid medium |  |  |  |
| Reagent | Supplier | Catalog Number | Final concentration |
| Basal medium |  |  | 40% |
| WNT CM | stable cell line |  | 25% |
| R-Spondin CM | stable cell line |  | 25% |
| Noggin CM | stable cell line |  | 10% |
| B27 supplement | Thermo Fisher Scientific | 17504044 | 1x |
| N2 supplement | Thermo Fisher Scientific | 17502048 | 1x |
| EGF | Peptrotech | AF-100-15 | 50 ng/mL |
| FGF-10 | Peptrotech | 100-26 | 100 ng/mL |
| Nicotinamide | Sigma | 3376 | 1 mM |
| TGF-β inhibitor | Tocris | 2939 | 0.5 uM |

## Supporting information captions

**S1 Fig. The morphology of uninfected GI organoids and monolayers.** Representative phase-contrast images of the human gastric (corporal), small intestinal (jejunal), large intestinal (colonic) organoids in a Matrigel drop and the subconfluent 2D monolayers derived from respective organoid cultures in microwell plates. Scale bar: 100 µm.

**S2 Fig. *C. trachomatis* infects Ki67-positive and negative gastric cells.** (A) Subconfluent gastric cells in microwell plates were infected with GFP-expressing *C. trachomatis* (MOI of 5) and 24 hours p.i. fixed, stained and subjected to confocal microscopy. Representative fluorescence microscopic images of three independent experiments show the localization of Ki67-positive cells in uninfected and infected samples (blue: DAPI, green: *C. trachomatis*, grey: actin, magenta: Ki67). Scale bar: 50 µm. (B) The percentage of the chlamydial inclusions residing in Ki67-positive or negative cells was determined by manually quantifying inclusions in five fields of view per sample. Data represent mean values ± SD from three independent experiments. Statistical analysis of the data was performed using unpaired t-test (ns = not significant).

**S3 Fig. The effect of EGTA treatment on the morphology and infection pattern of intestinal cells.** (A) Small and large intestinal epithelial cells grown as subconfluent monolayers were treated with 4 mM EGTA for 30 min at 37°C or left untreated. Phase-contrast images show the changes in the morphology of the cells upon treatment. Scale bar: 100 µm. (B) Intestinal cells pre-treated with 4 mM EGTA for 30 min or left untreated, were infected with GFP-expressing *C. trachomatis* (MOI of 5). 24 hours p.i. the cells were fixed, stained and subjected to confocal microscopy (blue: DAPI, green: *C. trachomatis*, red: Actin). Scale bar: 100 µm.

**S4 Fig. *C. trachomatis* apical versus basolateral infection in human fallopian tube cells.** (A) Organoid-derived human primary fallopian tube cells grown on cell culture inserts were infected with GFP-expressing *C. trachomatis* (MOI of 5) from apical or basolateral surface․ After 2 hours of incubation, the inoculum was removed and the cells were kept in the fresh organoid medium. 24 hours p.i. the cells were fixed, stained and subjected to confocal microscopy. Shown are representative confocal fluorescence Z-stack images of four independent experiments (blue: DAPI, green: *C. trachomatis*). Scale bar: 50 µm. (B) The numbers of chlamydial inclusions and host cell nuclei in (A) were quantified in at least 4 fields of view per sample using Fiji. Data represent mean values ± SD from four independent experiments. Statistical analysis was performed using an unpaired t-test (*P<0.05). The data point numbers and shapes below the graphs refer to the donor IDs used in the experiments.

**S5 Fig. Infection of the Pgp3-deficient and plasmid-free *C. trachomatis* in HeLa cells.** (A) Primary infection of HeLa cells infected with *Ctr* WT, *Ctr* Δ*pgp3* and *Ctr* PF. Subconfluent monolayers of HeLa cells were infected with the chlamydial derivatives at MOI of 0.5. 24 hours p.i. the cells were fixed, stained and the infection rate was determined by quantifying the numbers of inclusions and host cell nuclei in 14 fields of view per sample by automated microscopy. To assess the infectivity of the chlamydial progeny (B), the infected HeLa cells were lysed 48 hours p.i. and freshly seeded HeLa cells were infected with an aliquot of the lysates. 24 hours p.i. the cells were fixed, stained and the infection rate was determined in 14 fields of view per sample by automated microscopy. (C) The average size of the inclusions during primary infection was determined by automated microscopy in 14 fields of view per sample. All graphs represent mean values ± SD from three independent experiments. Statistical significance was determined by one-way ANOVA (ns = not significant, *P<0.05, **P<0.01, ***P<0.001).

